# Rapid generation of CD19 CAR-T Cells by minicircle DNA enables anti-tumor activity and prevents fatal CAR-B leukemia

**DOI:** 10.1101/2022.11.25.517980

**Authors:** Xueshuai Ye, Yongqiang Wu, Ziqi Cai, Qichen Yuan, Jianhui Cai

## Abstract

Manufacturing CAR (chimeric antigen receptor)-T cell with viral vector is expensive and time-consuming. Besides, during viral transduction, the genes encoding CARs are randomly integrated into the genome, which could cause oncogenesis or produce devastating CAR-tumor cells. Here, using virus-free and non-transgenic minicircle DNA (mcDNA) vector, we enable rapid generation of CD19 CAR-T cells within two days. Further, we demonstrate in vitro and in xenograft models that the anti-tumor effects of CD19 CAR-T cells produced by mcDNA are as effective as those produced by viral vectors. Finally, we show that our manufacturing process can avoid the production of fatal CAR-tumor cells. Taken together, we provide a fast, effective, and therapeutically safe method to generate CD19 CAR-T cells for treating leukemia.

## Introduction

Chimeric antigen receptors T cells therapy (CAR-T) is one of the most effective adoptive cell therapies (ACT) in treating hematological malignancy and many types of solid tumors. The CAR structure usually consists of three portions, including extracellular antigen recognition, transmembrane, and intracellular T cell activation domains(*1*). The extracellular portion of a CAR is a single-chain fragment variable (scFv) derived from a monoclonal antibody that can target tumor-specific antigens (TSA), such as CD19, CD20, or CD22(*2, 3*). The transmembrane portion is derived from CD28 or CD8 and regulates the molecular interactions associated with CAR signaling by forming homodimers or trimers(*4*). Lastly, the intracellular signaling domain, typically the zeta subunit of the CD3 complex, allows the CAR-expressing T cells to recognize the antigen on tumor cells independent of antigen processing and MHC presentation(*5*). So far, CAR-T therapy has been widely used in treating CD19 antigen-positive hematological malignancies, such as pediatric diffuse large B-cell lymphoma (DLBCL) and mantle cell lymphoma(*6*).

However, significant obstacles are still existing. First, the current clinical applications of CAR-T cells use lentiviral or retroviral vectors that randomly integrate the CAR gene into the human genome(*7*), causing insertional mutations that lead to oncogenesis(*8, 9*). In addition, the preparation of the CAR-T using viral vectors is expensive and time-consuming, limiting its broad applications(*10*). For example, the lentiviral vector-based conventional CAR-T manufacturing usually takes 2-3 weeks(*9*); the cost of FDA-approved CAR-T therapies ranges from $373,000 to $475,000(*11*). The fatal flaw is that once tumor cells sneak into the raw T cells during CAR-T manufacturing, CAR gene can be inserted into the genome of those tumor cells, generating CAR-expressing tumor cells, on which the tumor associated antigens are blocked by its own CAR. Due to such antigen blockage, CAR-tumor cells could escape the recognition and lysis by CAR-T cells, which can result in lethal consequences in therapeutic applications.(*12*). Although the sleeping beauty transposon has been used as a non-viral approach to CAR-T manufacturing, it still has genotoxic risk and cannot avoid deadly CAR-tumors(*13, 14*).

In this study, we tackled challenges of conventional CAR-T cell manufacture by using minicircle DNA (mcDNA)(*15*). Because mcDNA is a small non-integrating episomal vector which compared to viral vector can increase the in vivo transgene expression without causing transcriptional silencing(*15, 16*). We demonstrated that the mcDNA-based CD19-CAR-T cells can be prepared within 48 hours and the mcDNA CAR vector can be easily obtained by low-cost plasmid extraction. Furthermore, we showed that when treating CD19 positive leukemia both in vitro and in vivo, the mcDNA based CD19-CAR-T achieved a remarkable and continuous anti-leukemia effect compared to conventional viral-based CD19-CAR-T. Unlike viral vectors that cause CD19-blockage in leukemia due to continuous CAR expression, mcDNA approach achieved non-persisted CD19 blockage, of which the leukemia cells can be eradicated when the antigen re-expressed on the cell surface. In summary, our work established an effective approach to manufacturing CAR-T cells in a fast, safe, and cost-effective fashion.

## Materials and Methods

### Animals

All animal experiments were performed with approval of Hebei Medical University Animal Care and Use Committee, Hebei, China. All NOD scid gamma (NSG) mouse (female, 6-8 weeks old) used in this study were purchased from Jackson Laboratory and randomly assigned to experimental or control groups.

### Cell lines

Raji and K562 cell line were obtained from the American Type Culture Collection and cultured in RPMI-1640 medium containing 10% fetal bovine serum (FBS), and 1% penicillin– streptomycin. HEK293T cell line was cultured in standard DMEM supplemented with 10% FBS, 1% penicillin–streptomycin and 2 mM glutamine.

### Plasmid construction

The anti-CD19 single chain Fragment variable (scFv) was derived from the FMC63 mAb as previously published(*17*). The CAR construction was generated by fusing GM-CSF signal peptide, anti-CD19 scFv, CD8α transmembrane domain, and cytoplasmic domains of 4-1BB and CD3 zeta. The DNA fragments encoding CD19-CAR construction were synthesized and cloned into either the MC-Easy™ (SBI, USA) parental minicircle plasmid MN501A-1 to form the parental minicircle CD19-CAR plasmid (pMC-Easy-CD19-CAR) or the GV401 plasmid (GV401-CD19-CAR) for viral production.

### Cloning of minicircle CD19-CAR plasmid and lentivirus production

The mcDNA-CD19-CAR production procedures were performed as previously described(*18*). Briefly, the competent cells of engineering E. coli strain ZYCY10P3S2T (SBI) were transformed with pMC-Easy-CD19-CAR and cultured in LB medium containing 50 μg/mL kanamycin, with a rotation speed at 250 rpm at 30°C overnight. Then the L-arabinose were added to induce the expression of ΦC31 integrase (recombinase) and SceI endonuclease in E. coli strain ZYCY10P3S2T to generate the mcDNA-CD19-CAR by degrading the bacterial backbone. After induction culture for 5 h, The mcDNA-CD19-CAR were obtained and purified by using QIAGEN’s EndoFree Plasmid Maxi Kit. The lentivirus vectors were obtained as previously described(*13, 19*). HEK293T packaging cells were transfected with the CAR expressing plasmids (GV401-CD19-CAR) together with lentiviral packaging plasmid using Lipofectamine Reagent (Thermo Fisher Scientific). After 48 h, the culture supernatants containing lentivirus vectors (LV-CD19-CAR) were harvested and used for gene transduction.

### Preparation of human T cells

Peripheral blood samples were obtained from healthy donors at Hebei Blood Center, Hebei, China. PBMCs (Peripheral Blood Mononuclear Cells) were isolated from whole blood samples by Ficoll-Paque density gradient centrifugation (GE Healthcare) and collected, washed with complete medium for three times. For activation and transduction, CD8+ T cells were sorted from PBMCs using magnetic bead-based cell sorting CD8 MicroBeads (Miltenyi Biotec), the process was performed according to the manufacture’s instruction. The purity of sorted CD8+ T cells (from PBMC) was determined by flow cytometry. The sorted CD8+ cells were then cultured in RPMI-160 medium supplemented with 10% FBS and activated with anti-CD3 antibody (Thermo Fisher Scientific), anti-CD28 antibody (Thermo Fisher Scientific), 100 IU/ml interleukin-2 (IL-2, Novartis Proleukin). Cells were maintained at 37°C in a humidified atmosphere containing 5% CO_2_ for 2 days.

### Generation of MC-based CD19-CAR-T cells and conventional CD19-CAR-T

For generation of the conventional CD19-CAR-T, standard CAR-T cells for the clinical setting were produced except for several modifications(*20*): CD8+ T cells were stimulated with 1 ug/ml OKT-3/CD28 antibodies (Miltenyi Biotec) for 72 h, then transduced using lentivirus at a multiplicity of infection of 5. T cells were then expanded using 300 IU/ml IL-2, transduction is performed on day 2 and human AB serum is supplemented to the culture medium. For generation of the MC-based CD19-CAR-T, the activated CD8+ T cells were transduced with mcDNA-CD19-CAR by using preset program EO115 of 4D-Nucleofector™ system (Lonza). Electroporation was performed according to the manufacturer’s instruction. After electroporation, cells were resuspended in pre-warmed RPMI-1640 medium containing 10% FBS, 500 U/mL IL-2 and incubated in OKT-3/CD28-mAb-coated culture plates at 37°C in 5% CO_2_ to reactivate cells. To verify the feasibility and stability of the MC-based CD19-CAR-T manufacture, the transfection experiments were repeated for 3 times with cells obtained from 3 different donors. After lentivirus transduction for at least 72 h or electroporation for 48 h, the CAR expression efficiencies, and the intensity of CAR expression of transfected T cells were detected by flow cytometry using FITC-Labeled Human CD19 (20-291) Protein(ACRO biosystem, China).

### Generation of MC-based CD19-CAR-Raji and lentivirus CD19-CAR-Raji cells

For generation of the MC-based CD19-CAR-Raji, the Raji cells were transduced with mcDNA-CD19-CAR by using preset program DS104 of 4D-Nucleofector™ system (Lonza). Electroporation was performed according to the manufacturer’s instruction. After electroporation, cells were resuspended in pre-warmed RPMI-1640 medium containing 10% FBS. LV-CD19-CAR were used to transfect Raji at a multiplicity of infection of 5. After lentivirus transduction for at least 72 h or electroporation for 48 h, the human CD19 MicroBeads (Miltenyi Biotec) were used for depletion of CD19+ cells and the CD19 expression of transfected Raji cells were detected using PE-labeled anti-CD19 (Santa Cruz).

### Flow cytometry analysis

Raji and K562 were stained with a PE-labeled anti-CD19 (Santa Cruz) and analyzed by FACS for CD19 expression on the cell surface. The CD8+ T cell phenotypic and its rate change before and after the sorting process were analyzed by flow cytometry with CD8-FITC (eBioscience). The CD19 CAR expression on the surface of CAR-T cells was measured FITC-Labeled Human CD19 (20-291) Protein (ACRO biosystem, China).

### Cell proliferation assay

To study the correlation between the cell proliferation and the CAR-expression of MC-based CD19-CAR-T cells, 1×10^6^ MC-based CD19-CAR-T cells were processed as single-cell suspension and added with equal volume of CFSE working fluid (abmole) to label the viable cell. After incubated at 37°C for 10 min, labeling was terminated with 40% FBS immediately. After washing twice, the cells were resuspended with complete culture medium, the cell number and CAR expression were detected by flow cytometry every other day and the proliferation curve was generated.

### Cytotoxicity assay

Cytotoxic activities of either conventional CD19-CAR-T cells or MC-based CD19-CAR-T were assessed by Annexin V-Cy5 reagent (Biovision) according to manufacturer’s instructions. The Raji cells and K562 were set as target cells and seeded 4 × 10^5^ cells/well, respectively. Effector T cells were then added at 4 × 10^5^ cells/well and co-incubated at 37°C. After 4 h, cells were collected and stained for apoptosis by Annexin V-Cy5 reagent (Biovision). Target cells were assessed for AnnexinV staining by flow cytometry. The killing assays for MC-based CD19-CAR-Raji (day1, 7, 14, 21) and lentivirus CD19-CAR-Raji (day1, 7, 14, 21) were also performed as described above and the Raji cells were collected and stained for apoptosis by Annexin V-Cy5 reagent (Biovision).

### ELISA analysis for IFN-γ and IL-2 secretion

For IFN-γ and IL-2 secretion tests, effector cells (1 × 10^5^ cells/100 ul) and target cells (1 × 10^5^ cells/100 ul) were plated in 96-well plates. Supernatants were collected after co-culturing for 24 h and ELISA for the detection of human IFN-γ or IL-2 was carried out using Human IFN-γ testing ELISA kit or Human IL-2 testing ELISA kit (both from eBioscience).

### CD19-CAR-T treatment in vivo

To evaluate the cytotoxicity of CD19-CAR-T cells in vivo, a mixture of Raji cells (CD19+ GFP+) and K562 target cells (CD19-RFP+) were resuspended in PBS (1 × 10^7^ cells per cell type) and injected into the intraperitoneal space (i.p.) in each (NSG) mouse. 14 hours later, 2 × 10^7^ either conventional CD19-CAR-T cells or MC-based CD19-CAR-T cells were injected (i.p.) using the same administration route. Mice were euthanized using carbon dioxide asphyxiation followed by cervical dislocation. Peritoneal lavage was collected based on a procedure described previously(*21*). Briefly, 1.5 ml of cold PBS was used to re-suspend cells in the peritoneal cavity. The lavage was centrifuged at 400 g and room temperature for 5 min, re-suspended in 0.5 mL of red blood cell lysis solution at 1 x concentration (BD Biosciences) for 15 min at room temperature, centrifuged again at 400 g for 5 min, and fixed in PBS for FACS analysis.

To establish the Raji (CD19+) tumor model, each NSG mice were subcutaneously inoculated with 5 × 10^6^ Raji cells(*22*). 5 days after injection, tumor engraftment was evaluated by Vernier-caliper and the mice with comparable tumor loads were segregated into normal saline group (treated with 0.2 ml), conventional CD19-CAR-T cell group (treated with 2 × 10^7^ conventional CD19-CAR-T/0.2 ml), MC-CD19-CAR-T cell group (treated with 2 × 10^7^ MC-CD19-CAR-T/0.2 ml). The tumor size was measured using a Vernier-caliper every other day and the survival time of all the mice were monitored consistently. Tumor volume was calculated as: Volume = (length × width^2^) / 2.

### CD19-CAR-Raji treatment

For the CD19-CAR-Raji tumor model, the transfected Raji cells were sorted with human CD19 MicroBeads (Miltenyi Biotec) and each NSG mice were subcutaneously inoculated with either 5 × 10^6^ MC-CD19-CAR-Raji and lentivirus CD19-CAR-Raji. Tumor engraftment was evaluated by Vernier-caliper and the mice with comparable tumor loads were segregated into normal saline group (treated with 0.2 ml), conventional CD19-CAR-T cell group (treated with 2 × 10^7^ conventional CD19-CAR-T/0.2ml), MC-CD19-CAR-T cell group (treated with 2 × 10^7^ MC-CD19-CAR-T/0.2ml).The tumor size was measured using a Vernier-caliper every other day and the survival time of all the mice were monitored consistently. Tumor volume was calculated as: Volume = (length × width^2^) / 2.

### Statistical Analysis

All statistical analyses were performed using the Prism 9.3.1 (GraphPad Software). Statistical comparisons between two groups were determined by two-tailed parametric or nonparametric (Mann–Whitney U-test) t-tests for unpaired data or by two-tailed paired Student’ s t-tests for matched samples (produced from same donor). P < 0.05 were considered statistically significant.

## Results

### The mcDNA-CD19-CAR vector is efficiently expressed in human T cells and provides two weeks of transgene expression

Firstly, we cloned the anti-CD19 CAR construct by fusing GM-CSF signal peptide, anti-CD19 scFv, CD8α hinge and transmembrane domain, the 4-1BB cytoplasmic domains and CD3 zeta all together (Fig.1a). The mcDNA-CD19-CAR was generated in engineered bacteria ZYCY10P3S2T (Fig.1b, Fig.S1). To prepare the CAR-T cells, CD8+ T cells were obtained from peripheral blood and sorted by magnetic beads. FACS analysis demonstrated that the percentage of CD8+ cells in PBMCs ranges from 10%-20% and all the samples can be enriched up to nearly 100% (Fig.1c). 48 h after transfection, CAR expression can be detected by FACS analysis. The results showed that the transfection efficiency of mcDNA-CD19-CAR-T (CD19-CAR-T cells) was 73.57%, which is higher than the 43.70% of lentivirus transfected cells (conventional approach). Cells without lentivirus transfection or electroporation can hardly be detected via the FITC-positive signals (Fig.1d).

**Figure 1.**
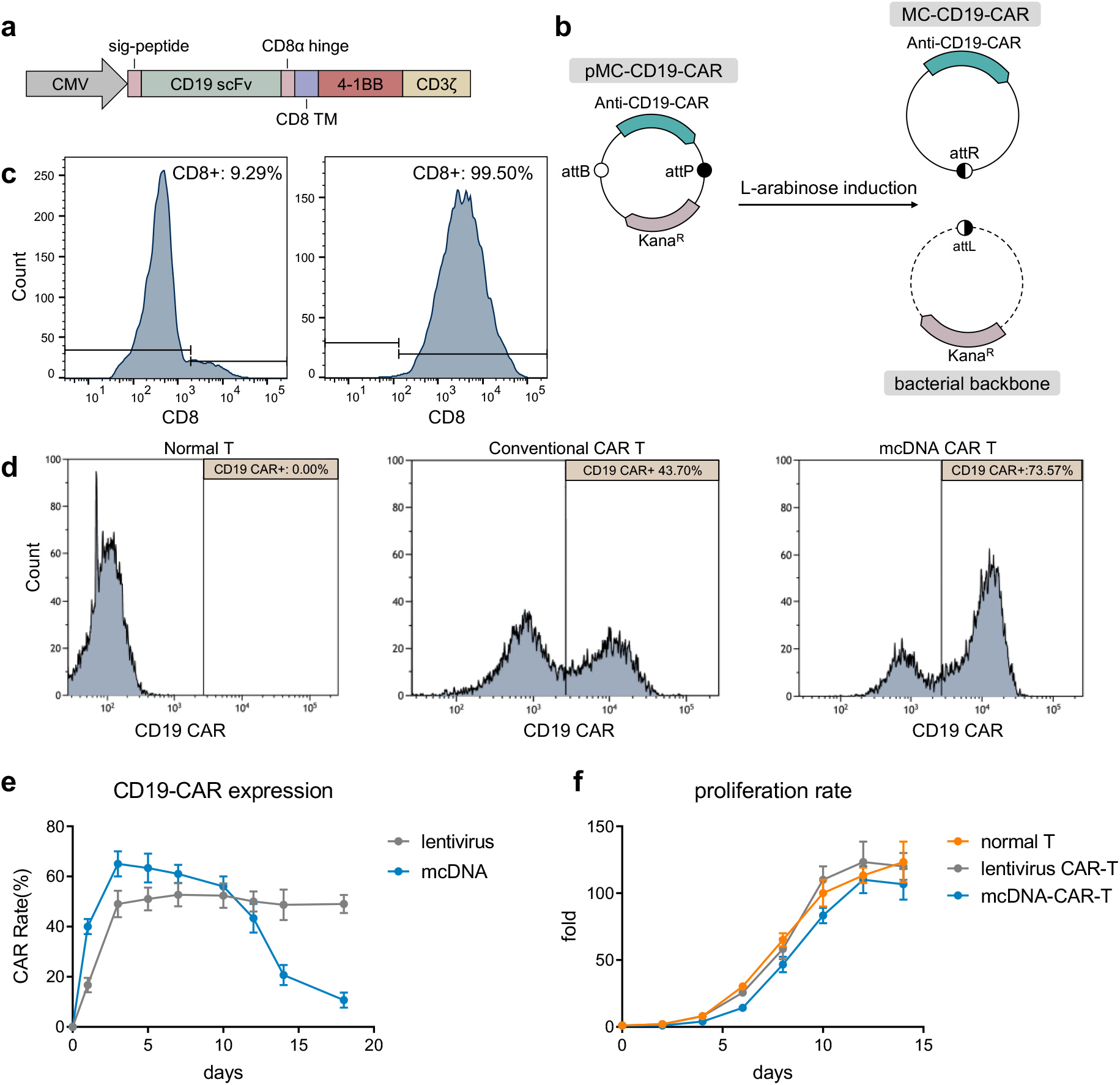
Generation of mcDNA-CD19-CAR-T cells. (a) Schematic of CD19 CAR structure. (b) Schematic of the process to generate CD19-CAR mcDNA in bacterial E.coli strain ZYCY10P3S2T. After L-arabinose induction, pMC-CD19-CAR was separated into mcDNA-CD19-CAR and the bacterial backbone was removed by SceI endonuclease digestion. (c) Human CD8+ T cells were sorted from PBMCs using MACS™. Samples of PBMCs were obtained from three different healthy donors and evaluated by FACS (left), after the sorting process, CD8+ ratio were evaluated by FACS (right). (d) 72 hours after electroporation or lentivirus transfection, the CAR expression of T cells was evaluated by FACS, and non-transfected normal T cells were used as a control (left). (e) Eighteen days after electroporation, CD19-CAR expression was consistently tested by FACS. (f) Proliferation curves of normal T, lentivirus CAR-T, mcDNA-CAR-T cells. The statistical analysis was performed on cells from 3 donors.

To detect the persistence time and the amount of CAR expression after electroporation of T cells. The CAR expression level was monitored routinely for the percentage of CAR+ cells (CAR %). Based on the statistical analysis from 3 donors, we observed that mcDNA-CD19-CAR-T cells reached the highest expression of CD19-CAR on day 3 (Fig.1e). We show that as the mcDNA-CD19-CAR-T cells cultured from day 1 to day 14, the CAR % dropped from 70% to 20% (Fig.1e). Meanwhile, stable expression of CAR was observed in lentivirus transfected T cells (Fig.1e).

To investigate the impact of the mcDNA vectors on primary human T cells, the isolated CD8+ cells manufactured with mcDNA were compared with T cells transduced with lentivirus carrying the same expression cassette. Although slow at the beginning, the proliferation rate of mcDNA-CAR-T cells increased to the similar levels as lentivirus transfected T cells and normal T cells after 10 days (Fig.1f).

### Rapid MC-CD19-CAR-T cells specifically lyse CD19 positive leukemia cells both in vitro and in vivo

To evaluate in vitro anti-tumor reactivity and specificity of rapid manufactured MC-CD19-CAR-T, we chose human leukemia cell lines Raji (CD19+) and K562 (CD19-) for the test. We confirmed that Raji over-expresses CD19 antigen and the K562 cells are CD19-negative (Fig.2a). Next, we performed the in vitro killing assays by co-incubating the conventional CAR-T or different stages of MC-CD19-CAR-T with target cells according to previous methods(*23*), in which normal T cells, conventional CD19-CAR-T cells, mcDNA-CD19-CAR-T cells (1-day, 6-days, 13-days post-electroporation) were co-cultured with Raji and K562 mixture for 4 h at two-to-one ratio of effector-to-target cells, respectively. FACS analysis showed that conventional CD19-CAR-T exhibited strong cytotoxicity against CD19-positive Raji cells but not against CD19-negative K562 cells. The mcDNA-CD19-CAR-T cells 1-day post-electroporation (mcDNA-CD19-CAR-T D2) and 6-days post-electroporation (mcDNA-CD19-CAR-T D7) showed similar trend as conventional CD19-CAR-T, while moderate killing activity was observed using mcDNA-CD19-CAR-T D14 (13-days post-electroporation), indicating that the CAR expression levels of mcDNA-CD19-CAR-T cells decreased over time after electroporation (Fig. 2b). In addition, normal T cells showed no significant cytotoxicity against either CD19-positive Raji cells or CD19-negative K562 cells, indicating specific killing mediated only by CD19-CAR-T cells (Fig. 2b).

**Figure 2.**
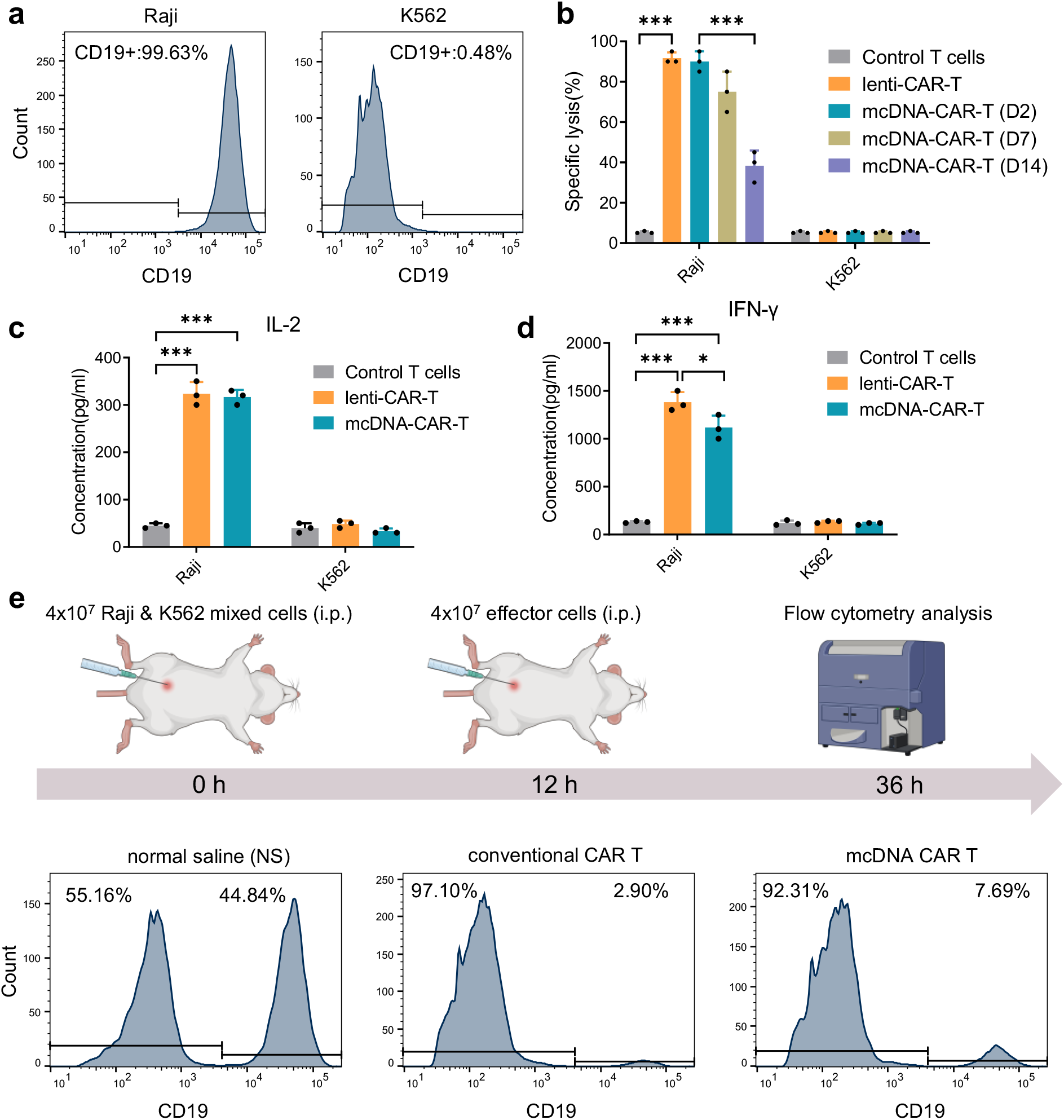
The mcDNA-CAR-T cells can specifically and efficiently target CD19-positive cells in vitro and in vivo. (a) FACS analysis of the surface expression of CD19 on Raji and K562 cell lines. (b) Histograms showing the CD19-specific cytotoxicity against Raji cells at the ratio of 2:1 (effector to target cell ratio). mcDNA-CAR-T cells were obtained and tested from 3 donors (1, 7, 14 days after electroporation, respectively). Statistical analysis was performed by nonparametric test. (c) ELISA analysis of cytokine production of IFN-γ in the co-culture medium. (d) ELISA analysis of cytokine production of IL-2 in the co-culture mediums. Statistical analysis was performed by nonparametric tests. (e) Schematic of the mouse model used and the representative results of tumor cell survival. Matched CD19+ and CD19-target cells were injected into the intraperitoneal space of NOD scid gamma (NSG) immune-deficient mice. Normal saline (NS), conventional CAR-T cells, or mcDNA-CAR-T cells were injected (i.p.) at the indicated times. At the end of the experiment, flow cytometry was performed to measure the ratio of surviving CD19+ versus CD19-target cells.

To detect the cytokine secretion of either conventional CD19-CAR-T or MC-DNA-CAR-T cells, we performed ELISA assay and showed that conventional CD19-CAR-T and mcDNA-CAR-T cells significantly increased the IFN-γ or IL-2 secretion when co-cultured with CD19-positive Raji cells, while only little amount of IFN-γor IL-2 can be detected when co-cultured with CD19-negative K562 cells (Figs. 2c and 2d). These results demonstrate that rapid mcDNA-CD19-CAR-T can specifically and effectively lyse CD19 positive leukemia cells. To evaluate the short-term and specific cytotoxic of mcDNA-CAR-T in vivo, we established a hematologic malignancy mouse xenograft model using the mixture of Raji (CD19+) and K562 (CD19-). The results show that both conventional CAR-T and mcDNA-CD19-CAR-T can lyse the CD19+ Raji cells population specifically in vivo (Fig. 2e).

### The mcDNA-CAR-T manufacturing prevents the survival of CAR-tumor cells in vitro

Because mcDNA-CAR-T cells can limit the expression of CAR, which might prevent the survival of deadly CAR-tumor cells that are unexpectedly generated during the manufacturing. To demonstrate this potential advantage of mcDNA-CAR-T over conventional CAR-T manufacturing, we used CD19 positive Raji cells as a tumor cell model to produce CAR-tumor cells. We transfected or transduced the Raji cells with mcDNA-CD19-CAR vector or lentivirus-CD19 CAR vector and sorted the CD19 antigen undetectable Raji cells (CAR-Raji cells). Flow cytometry analysis showed that the Magnetic Activated Cell Sorting (MACS) procedure can successfully enrich both lentivirus-transduced and mcDNA-transfected CAR-Raji cells (Fig. 3a). We then tested the cytotoxicity of CD19 CAR-T cells against CAR-Raji cells (target cells) by time course using conventional CD19 CAR-T cells as effector cells with a ratio of 2:1 (effector to target cell). We performed five tests within three weeks but did not observe lysis of lenti-CAR-Raji cells. However, the lysis of mcDNA-CAR-Raji increased over time, reaching 82% by day 21 (Fig. 3b). We used FACS to quantify CD19 antigen of lenti-CAR Raji and mcDNA-CAR-Raji, tested five times within four weeks, in which we did not observe CD19 antigen of lenti-CAR-Raji, but the CD19 antigen of mcDNA-CAR-Raji elevated gradually and achieved up to 92 % after four weeks (Fig. 3c).

**Figure 3.**
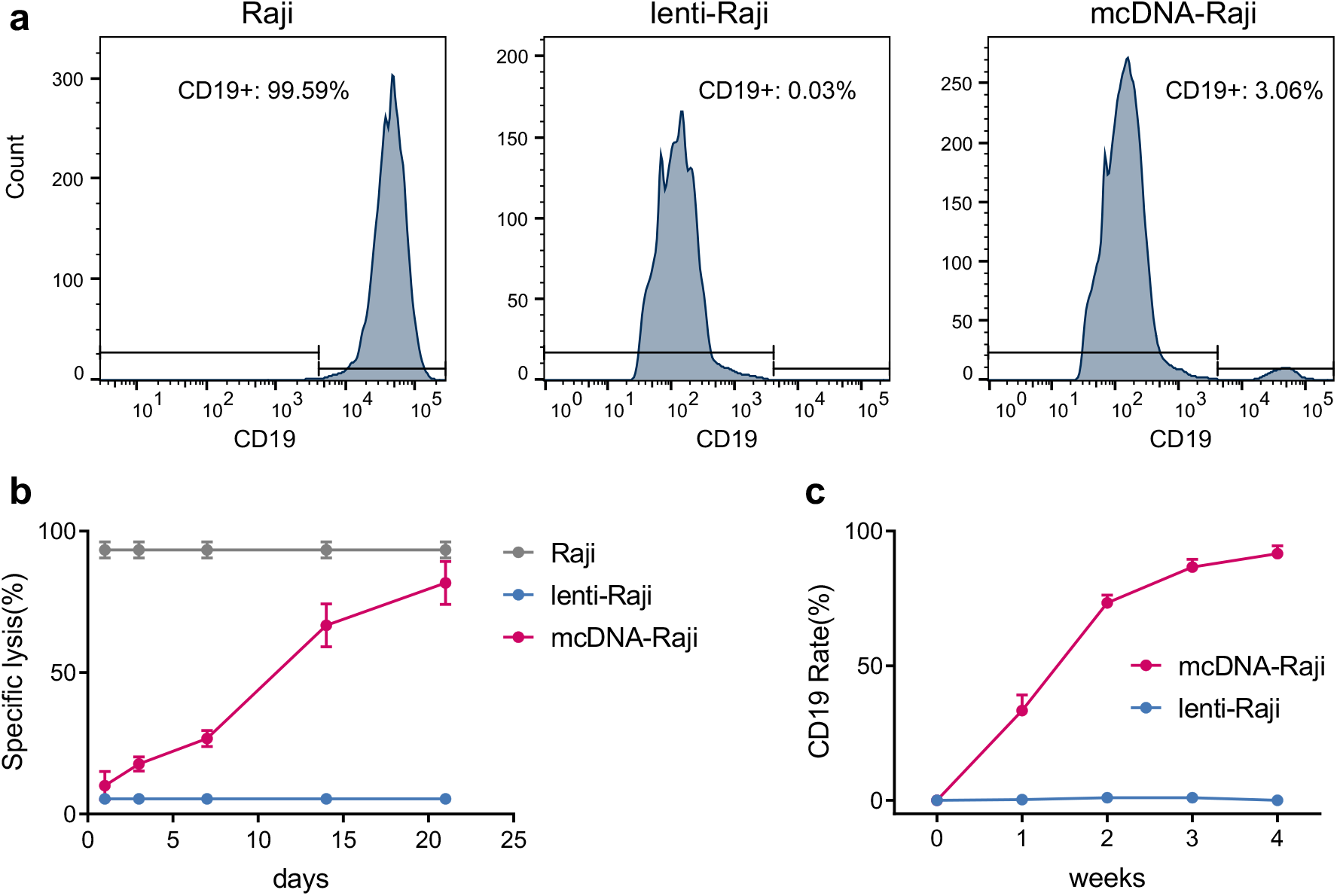
CD19-CAR blocks the CD19 antigen on the Raji cell. (a) FACS analysis of the surface expression of CD19 on mcDNA-Raji and lenti-Raji cell lines. (b) Histograms showing the CD19-specific cytotoxicity against Raji cells at the ratio of 2:1 (effector to target cell ratio). mcDNA-CAR-T cells were obtained and tested from 3 donors (1, 3, 7, 14, 21 days after electroporation, respectively). (c) CD19 expression periods of mcDNA-Raji and lenti-Raji. CD19 antigen on Raji cells transfected with mcDNA could be blocked but reappeared 3 weeks after transfection, while Raji cells transfected with lentivirus did not show this trend.

### The mcDNA-CAR-T manufacturing prevents fatal CAR-tumor cells in xenografted mouse model

To test the long-term tumor-killing effect of mcDNA-CD19-CAR-T cells, we established a tumor model by subcutaneously implanting Raji (CD19+) cells into NSG mice. On day 5 after tumor loading, mice with tumors reaching 100-150 mm^3^ were randomly assigned to four test groups, three of which received single-dose tail vein injection with normal saline, mcDNA-CD19-CAR-T, and conventional CAR-T, respectively. One group was administrated two-dose tail vein injections of mcDNA-CD19-CAR-T cells on day 5 and 13 after tumor loading (Fig. 4a). We found that a single dose of mcDNA-CD19 CAR-T cells can inhibit tumor growth for up to ten days but failed to continue being effective, compared to conventional CAR-T cells and normal saline groups (Fig. 4b and 4c). However, two doses of mcDNA-CD19 CAR-T achieved the same tumor-killing efficacy as using conventional CAR-T (Fig. 4b and 4c), suggesting effective therapeutic treatment with multiple doses of mcDNA-CD19-CAR-T.

**Figure 4.**
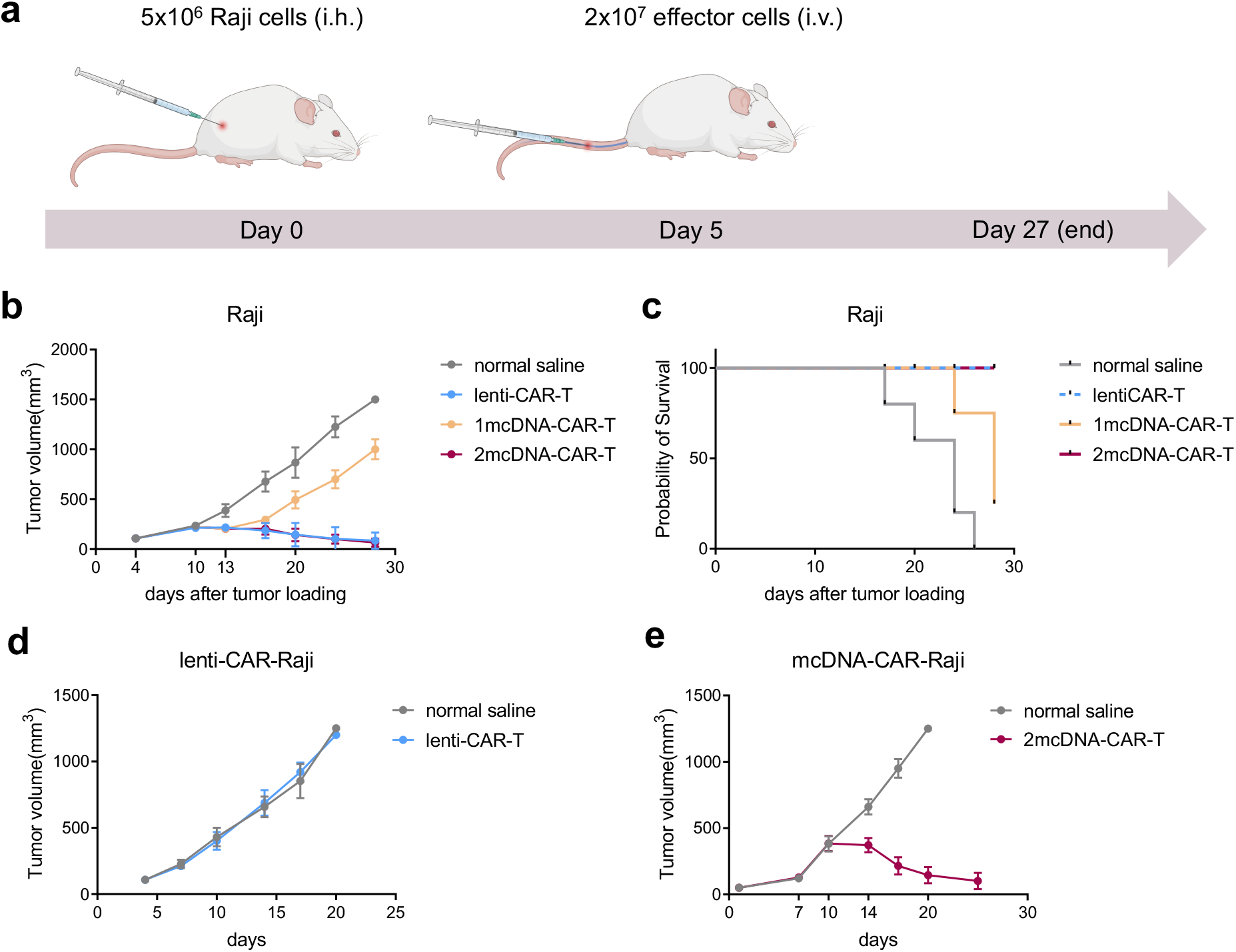
In vivo lysis of mcDNA-CAR-Raji by CD19-CAR-T cells. (a) Schematic of the mouse models bearing either Raji, mcDNA-CAR-Raji or lenti-CAR-Raji xenograft tumor model. (b) Tumor volume of Raji xenograft mice. When the tumor volume reached around 100 mm^3^, 1×10^7^ effector cells were injected on day 4 and 13. 1mcDNA-CAR-T, injected on day 4; 2mcDNA-CAR-T, injected on both day 4 and 13. The volumes of xenograft mice tumors were recorded for statistical analysis (statistics presented as mean ± SD, n = 5 for each group). (c) The percent survival curve of each Raji xenograft mice groups. (e) Tumor volume of lenti-CAR-Raji xenograft mice. (f) Tumor volume of mc-CAR-Raji xenograft mice. The treatment was performed in Raji xenograft mice group. 2mcDNA-CAR-T, injected on both day 7 and 14.

To evaluate the tumorigenicity of CAR-Raji cells and their response to CAR-T therapy in vivo, we established xenografted mouse models using CD19-CAR-Raji produced by either mcDNA or lentivirus, respectively. Then we sought to test the cytotoxicity of mcDNA-CAR-T and conventional CAR-T cells in Raji, mcDNA-CD19-CAR Raji and lenti-CD19-CAR Raji xenograft tumor models (Fig. 4a). In the lentivirus-CD19-CAR-Raji mouse model, no significant difference of tumor size was observed between conventional CAR-T and normal saline treated groups (Fig. 4d), and the mice received the conventional CAR-T did not show survival advantages over normal saline treated mice (Supplementary Fig. 2a). However, the mcDNA-CD19-CAR-T treatment substantially decreased the tumor size of mcDNA-CD19-CAR Raji xenografted mouse models (Fig. 4e), resulting in prolonged survival in comparison to the groups treated with normal saline (Fig. 4e, Supplementary Fig. 2d). Notably, the growth of mcDNA-CD19-CAR-Raji cells, after treated with a single dose of the mcDNA-CD19-CAR-T cells, were not effectively inhibited until 9 days after tumor loading; however, using a second dose of mcDNA-CD19-CAR-T cells, significant tumor regression was observed 14 days after tumor loading (Supplementary Figs. 2b and 2c). These results demonstrate that Raji cells modified with lentivirus-CD19 CAR can block the CD19 antigen permanently and cause deadly target loss. The mcDNA vector enables highly efficient manufacturing of the CAR-T cells without eliciting deadly antigen-blockage. For potential therapeutic applications, we suggest using multiple-dose of mcDNA-CD19-CAR-T cells to achieve effective inhibition of the tumor growth.

## Discussion

Here, using mcDNA, we demonstrate the rapid generation of CD19-CAR-T cells to prevent fatal CAR-B leukemia while maintaining the anti-tumor efficacy. We demonstrate that mcDNA-CD19-CAR-T cells can be produced efficiently, which provide at least 14 days of CAR expression. We show that mcDNA-CD19-CAR-T can specifically lyse CD19 positive leukemia cells both in vitro and in vivo. Since mcDNA is a non-integration gene delivery vector, the CD19 antigen blocking effect in mcDNA-CD19-CAR-Raji cells can be eliminated, and mcDNA-CD19-CAR-Raji can be lysed by multiple infusions of mcDNA-CD19-CAR-T cells. As a result, the mcDNA CAR-T production process can prevent the persistent CD19 antigen blockage, which can be caused by the virial vectors. Together, we demonstrate that mcDNA is a safe and efficient gene delivery vector for CAR-T manufacturing.

CAR-T cells have shown anti-tumor activity in a variety of preclinical and clinical studies(*24–26*), several CD19 targeted CAR-T products, such as Yescarta, Kymriah and Tecartus, are approved by the FDA for the treatment of lymphomas(*27*). However, all those approved CAR-T products are produced by viral vectors that can cause potential mutagenesis by random genomic integration of the CAR-coding genes(*28*). Furthermore, the manufacture of viral vectors is time-consuming and expensive, in which B lymphocytes cannot be completely removed from raw T cells, resulting in persistent CAR expression on virus-infected B cells, which can lead to consistent antigen blockade with fatal consequences. Similarly, the self-replicating non-integrating DNA vector can lead to consistent antigen blocking through consistent expression of CAR (*29*).

In our study, we demonstrate a convenient and safe method to generate CAR-T cells using mcDNA. Compared with viral vectors, mcDNA vectors are non-integrating, non-self-replicating, enabling longer but not persistent CAR expression. The mcDNA method does not cause continuous antigen blocking, and even when B lymphocytes are transfected with mcDNA-CAR, CAR-B cells can be lysed by the next infusion of CAR-T cells.

In summary, we demonstrate the rapid generation of CD19-CAR-T using mcDNA vectors, with manufacturing time compressed to within 48 hours, and without the risk of potentially lethal CAR tumors. The mcDNA CD19-CAR-T showed almost the same cytotoxicity as conventional CD19-CAR-T in vitro, and a durable anti-tumor ability could be achieved through multiple doses of mcDNA CD19-CAR-T infusion. Different CAR structure designs may affect the anti-tumor ability of mcDNA CAR-T, and we anticipate future studies to reveal the appropriate CAR structure and the expression intensity of mcDNA CAR on T cells.

## Declarations

## Acknowledgement

We thank Wentao Zhang, Yang Li for providing technical assistance. This work is supported by the Hebei government special project of top talents.

## Data Availability

Data are available from the corresponding authors upon reasonable request.

## Author contribution

Xueshuai Ye and Jianhui Cai initiated and designed the in vitro and in vivo studies. Xueshuai Ye and Yongqiang Wu performed the preclinical animal studies. Yongqiang Wu assisted in the construction of the vectors. Ziqi Cai assisted in managing the research project. Xueshuai Ye, Yongqiang Wu, Jianhui Cai prepared the manuscript draft. Qichen Yuan provided feedbacks, revised the figures, and wrote the manuscript. All authors read and approved the final manuscript.

## Ethics approval and consent to participate

The animals’ experimental protocol was conducted in accordance with the recommendations of the Guide for Care and Use of Laboratory Animals with respect to restraint, husbandry, surgical procedures, feed and fluid regulation, and veterinary care. All protocols were approved by Hebei medical university.

**Supplement Figure 1.**
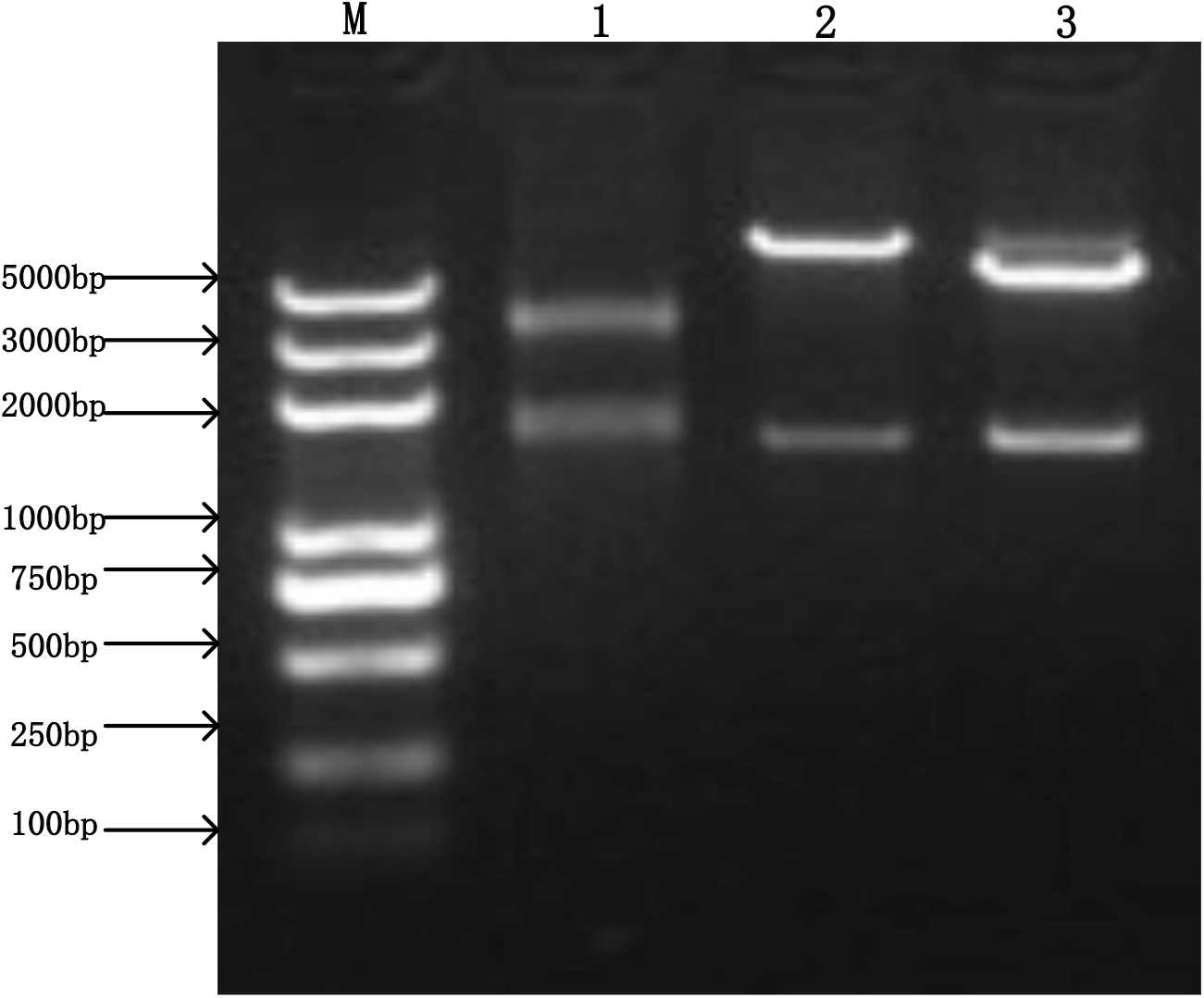
Electrophoresis for the closed circular plasmid and minicircle DNA before (−) and after (+) L-arabinose induction. M = molecular weight marker. Lane1: MC-CD19-CAR plasmid restriction enzyme digestion. Lane2: PLVX-CD19 CAR plasmid restriction enzyme digestion. Lane3: pMC-CD19-CAR plasmid restriction enzyme digestion.

**Supplement Figure 2.**
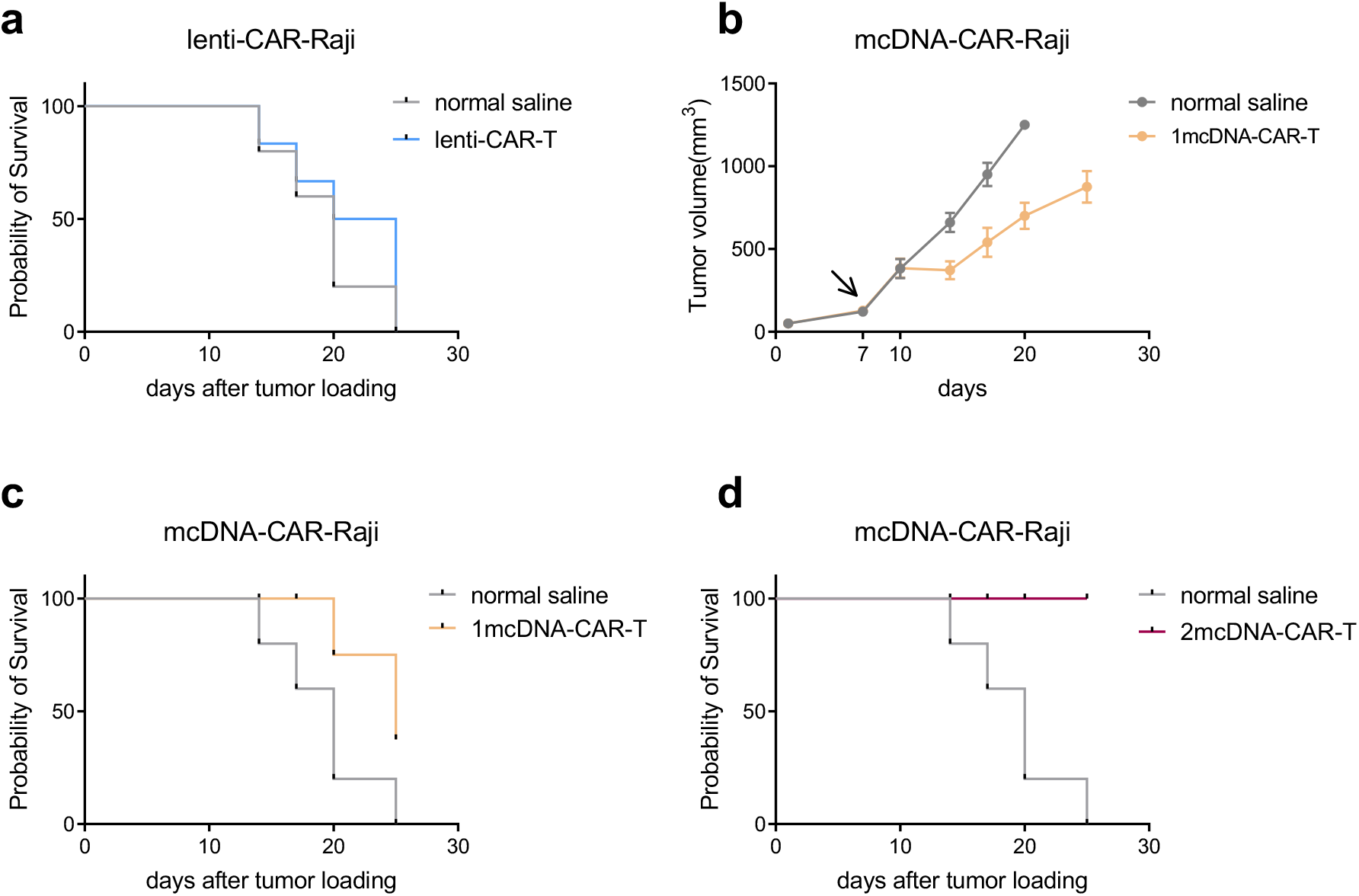
Percent survival and tumor volume of CAR-Raji xenografts. (a) Percent survival curve of Lenti-CAR-Raji xenograft mice groups. (b) Tumor volume of mc-CAR-Raji xenograft mice. The mice were given for single dose of mcDNA-CAR-T on day 7, as indicated by the arrow. (c) The percent survival curve of mcDNA-CAR-Raji xenograft mice groups treated with either single dose of mcDNA-CAR-T or normal saline. (d) the percent survival curve of mcDNA-CAR-Raji xenograft mice group treated with either two doses of mcDNA-CAR-T or normal saline.

